# Effect of OASL on oxaliplatin-induced immunogenic cell death in gastric cancer via the cGAS-STING signaling pathway

**DOI:** 10.1101/2024.12.09.627497

**Authors:** Lingling Zhang, Yi Liu, Haiying Yang, Luguang Liu, Longgang Wang, Jie Chai, Weizhu Zhao, Dong Sun

## Abstract

This study investigates the role of 2’-5’ oligoadenylate synthetase-like (OASL) in Oxaliplatin (OXA)-induced immunogenic cell death (ICD) in Gastric cancer (GC) cells through the cGAS-STING signaling pathway. Knockdown of OASL enhanced ICD expression, while overexpression had the opposite effect. RNA sequencing of OASL-knockdown and control GC cells treated with OXA revealed significant enrichment of the second messenger signaling pathway (cGAMP). cGAMP could directly activate STING as a second messenger, and cGAS was a key synthetic enzyme upstream of cGAMP. Next, the role of OASL in OXA-induced ICD in GC cells was validated through the cGAS-STING signaling pathway. The Co-IP and immunofluorescence results confirmed that the OASL and cGAS proteins can bind directly. Further research validated these findings in vivo. Results show that OASL regulates OXA-induced ICD in GC cells via the cGAS-STING pathway, impacting chemosensitivity. The findings suggest new targets and strategies for improving GC therapy by modulating OASL expression to enhance OXA sensitivity through immunogenic mechanisms.

## Introduction

Gastric cancer (GC) is the fifth most common malignancy worldwide, ranking as the fourth leading cause of cancer-related deaths, and is one of the most common malignant tumors in China ^[1,2]^. Although some progress has been made in the basic research and clinical application of GC in recent years, new chemotherapy drugs are constantly being applied in clinical practice, new therapeutic methods such as tumor immunotherapy and targeted therapy are constantly being updated, and the prognosis of GC patients remains poor ^[3–6]^. Chemotherapy plays an indispensable role in advanced GC^[7]^. Although Oxaliplatin(OXA)-based chemotherapy regimens can significantly improve the prognosis of patients with GC, the clinical response rate is only approximately 40–67% ^[8,9]^. Moreover, in recent years, it has been found that OXA is often prone to severe drug resistance in clinical medication, and previous studies have shown that changing the expression of some molecules in GC cells can improve their sensitivity of GC cells to OXA chemotherapy. However, these indicators have not yet been widely used in clinical practice ^[10,11]^. Therefore, it is important to conduct basic GC research at multiple levels to further clarify the molecular mechanisms involved in the development of GC.

2 ′ −5 ′ oligoadenylate synthetase (OAS) gene is one of the interferon-stimulated genes, OAS, a family of four members: OAS1, OAS2, OAS3, and ubiquitin-like 2 ′ −5 ′ oligo-adenylate synthetase (OASL), since OASL has an oligoadenylate-like domain at its N terminus, but the lack of oligoadenylate synthetase activity, with two ubiquitin-like domains at its C terminus, is a critical region for OASL to exert its antiviral activity^[12–15]^. Current studies have found that OASL plays an important role in immune regulation and maintenance of drug metabolism ^[16–18]^. A recent study found that the correlation between OASL and cancer has been gradually confirmed, which may participate in the development of breast cancer ^[19]^, pancreatic cancer ^[20]^, and cervical cancer ^[21]^, and is closely related to prognosis. Our previous study also found that OASL promotes the proliferation, invasion, and migration of GC cells and inhibits their apoptosis, and that OASL functional as an oncogene in the development of GC^[22]^. OASL is a decisive regulator in maintaining the sensitivity of lung cancer cells to acRoots, which may be related to the development of drug resistance. Modulation of OASL could be an alternative strategy for improving drug efficacy during cancer treatment ^[23]^. OAS family members have all been identified as central genes in trastuzumab-resistant GC and controls^[24]^, suggesting that OASL also plays an important role in maintaining sensitivity to drug therapy in GC. Recent studies have shown that OXA plays an important role in addition to cytotoxicity and that its anti-tumor activity is closely related to the regulation and improvement of immune function^[25]^. The specific immunological mechanisms of anti-tumor activity are: the induction of immunogenic cell death (ICD), the effect on the STAT protein signaling pathway, and regulation of the tumor microenvironment^[26–30]^. Accordingly, we speculated that OASL may play an important role in maintaining OXA chemotherapy sensitivity in GC.

Therefore, this study began with the anti-tumor immunological mechanism of OXA. GC cells were treated with OXA after OASL knockdown or overexpression. ICD expression was enhanced after OASL knockdown, whereas OASL overexpression had the opposite effect. OASL knockdown of GC cells and common GC cells were administered OXA for 24h for mRNA sequencing, which significantly enriched the second messenger signaling pathway (cGAMP). This also indicates that OASL can regulate OXA-induced ICD in GC cells through the cGAS-STING signaling pathway, thus affecting chemosensitivity. The results of this study provide new targets and strategies for GC therapy.

## Materials and methods

### 2.1 Cell culture

STAD cell lines (AGS, MKN45, and HGC27) were purchased from Cobioer (Nanjing, China). AGS,MKN45, and HGC27 cells were cultured in RPMI 1640 (BasalMedia, Shanghai, China) containing 10% fetal bovine serum (FBS) in a humidified incubator with 5% CO2 at 37°C.

### 2.2 Cell transfection

siRNAs (si-OASL) and the negative control (si-NC) were purchased from RiboBio (Guangzhou, China). OASL-amplified products were ligated into the pc-DNA3.1 expression vector (Tsingke Biotechnology, Beijing, China). si-OASL and si-NC were transfected into AGS and MKN45 cells, respectively, using Lipofectamine 2000 (Invitrogen,USA). The empty vector (Vector) and pcDNA3.1-OASL were transfected into HGC27 cells using Lipofectamine 2000. After 48 h, the transfection efficiency was measured by real-time quantitative polymerase chain reaction (RT-qPCR) and western blotting.

### 2.3 Cell counting kit-8 (CCK-8)

AGS, MKN45, and HGC27 cells were seeded in 96 well plates (5×l0^3^ cells/well). After incubation at the specified time, the CCK-8 solution (Beyotime, Shanghai,China) was added and incubated for another 2 h. The OD450 values were measured using a microplate reader (Bio-Rad, USA).

### 2.4 Flow cytometry

AGS, MKN45, and HGC27 cells were washed with PBS and digested with trypsin (0.25%). The cells were collected after centrifugation at 1000g for 5 min, suspended in PBS, and counted. Subsequently, the cells (2×10^5^ cells/well) were centrifuged and resuspended in 195 μL of Annexin V-FITC binding solution. Annexin V-FITC (5 μL) and propidium iodide (PI) staining solution (10 μL) were added and mixed gently. After incubation in the dark for 20 min at 37°C, apoptosis was assessed by flow cytometry.

### 2.5 Western blot

Total protein was extracted from the tissues and cells using RIPA lysis buffer (Beyotime, Shanghai,China), and the protein concentration was detected using the BCA Protein Assay kit (Beyotime, Shanghai,China). Protein samples (30 μg) were separated using 10% SDS-PAGE and transferred onto PVDF membranes. The membranes were incubated with primary.The antibodies were incubated overnight at 4°C. Thereafter, the membranes were incubated with HRP-conjugated secondary antibody (ab205718, Abcam, USA) for 2 h at room temperature.Subsequently, the bands were visualized using an ECL kit (Beyotime, Shanghai,China) and quantified using ImageJ software.The following primary antibodies were used for the western blot analysis: OASL (ab229136, Abcam, UK), cGAS (ab224144, Abcam, UK), STING (ab252560, Abcam, UK),IRF3 (ab245341, Abcam, UK), CRT (ab227444, Abcam, UK),HSP90 (ab203126, Abcam, UK),HSP70 (ab194360, Abcam, UK),CD8 (ab316778, Abcam, UK),GAPDH (10494-1-AP, Proteintech, USA).

### 2.6 Co-IP

Cells were collected 48 h after transfection and gently rinsed twice with precooled 1 × PBS. Freshly prepared cracking working solution was added, cracked on ice for 30 min, transferred to an EP tube, and sonicated on ice for 3 min. The supernatant was centrifuged and transferred to a new EP tube. Protein A/G PLUS Agarose (15 μL) (Beyotime, Shanghai,China) was added to each tube and incubated at 4°C for 1 h. The supernatant was centrifuged and transferred to a new EP tube. Lysis buffer (60 μl was added to 15 μl of 5 × loading buffer and boiled at 1000 W for 10 min. antibody (1.5 μg/tube) was added to the remaining lysate and incubated at 4°C in the suspension apparatus for 1 h. Protein A/G PLUS agarose (40 μl/tube) was added to a mixture of antibody and lysis buffer and incubated overnight at 4°C in a suspension apparatus. The beads were washed four times with lysis solution. After suction, 40 μl of lysis buffer was added to each tube, 40 μl of 2 × loading buffer was added, and the sample was boiled at 4°C for 10 min. Western blotting analysis and detection were performed.

### 2.7 ELISA

The cell suspensions of each group were centrifuged, the cell supernatant was collected, and the instructions of the corresponding ELISA kit (Beyotime, Shanghai,China) were followed to sequentially detect the levels of HMGB1 in the supernatant of the group cells.

### 2.8 ATP

The cell supernatant was collected after processing, and an ATP colorimetric detection kit (Beyotime, Shanghai,China) was used to measure ATP levels in the supernatant of the group cells.

### 2.9 LDH

The cell supernatant was collected after processing the cells, and an LDH assay kit (Beyotime, Shanghai,China) was used to detect the LDH levels in the supernatant of the group cells.

### 2.10 Immunofluorescence

Take cells treated with each group, fix them with 4% paraformaldehyde for 10 minutes, wash them with PBS, and then use 0 5% Triton X-100 was passaged at room temperature for 20 minutes, then blocked and added with primary antibody. The cells were incubated at room temperature for 1 h, followed by addition of labeled secondary antibodies. Cells were counterstained with DAPI, observed under a fluorescence microscope, and photographed.

### 2.11 Mouse Tumor-bearing Mode

C57/BL6 mice (n=20, male, 6-8 weeks old) were purchased from Jinan Pengyue Experimental Animal Breeding Co. Ltd. SPF mouse-specific feed and high-pressure sterilized water were used for feeding in the experimental animal center.

1. Mice were kept in a relatively sterile constant-temperature environment for one week and randomly divided into four groups: sh-NC (n = 5), sh-OASL (n = 5), OXA + sh-NC (n = 5), and OXA + sh-OASL (n=5).
2. The transfected MFC cells were adjusted for cell density to 1.5×10^6^ /mL, and a 0.2 mL cell suspension was seeded in the right armpit of mice. A tumor diameter of 0.5 cm, it indicated successful molding.
3. The dose was calculated using a human-to-mouse surface area ratio of 0.002:6, and oxaliplatin was 10 mg/kg was 0.2 mL (1 time / 2 d).
4. Tumor volume was measured every three days after injection (volume = length and width×2/2). Mouse was killed 15 days later, and the tumor bodies were dissected, weighed, and photographed.
5. The tumor tissue was divided into three parts, with a portion of fresh tissue extracted for protein for western blotting, another portion of fresh tissue subjected to TUNEL staining, and the last part of fresh tissue dehydrated and subjected to CD8 immunohistochemical staining.

### 2.12 Statistical analysis

GraphPadPrism17 was used for statistical analyses, and data are expressed as mean ± standard deviation (SD).The Student’s t-test was used to analyze the differences between the two groups. *P* < 0.05 was considered statistically significant. Three replicates were used for each experiment.

## Results

### 1 Oxaliplatin is able to induce immunogenic cell death in gastric cancer cells

#### 1.1 Effect of oxaliplatin on the proliferation and apoptosis of gastric cancer cells

To explore the effect of OXA on the proliferative activity of GC cells, we treated AGS, MKN45, and HGC27 cells with different concentrations (0,5,10,10,20,40,80, and 160 μM) for 24,48,and 72h. Cell proliferation assays using the CCK-8 assay showed that OXA significantly inhibited the growth of GC cells in a time-and dose-dependent manner (Figure 1A). We concluded that OXA inhibited the proliferation of GC cells to varying degrees. Similarly, to explore the effect of OXA on apoptosis in GC cells, AGS, MKN45, and HGC27 cells with OXA at different concentrations (0,10,20, and 40 μM) for 48 h. Flow cytometry (Annexin-FITC/PI double staining) showed that the apoptosis rate was significantly increased, along with the increased OXA concentration, compared to the controls (*P* <0.001, Figure 1B). This indicates that OXA induced apoptosis in GC cells in a dose-dependent manner.

**Figure 1.**
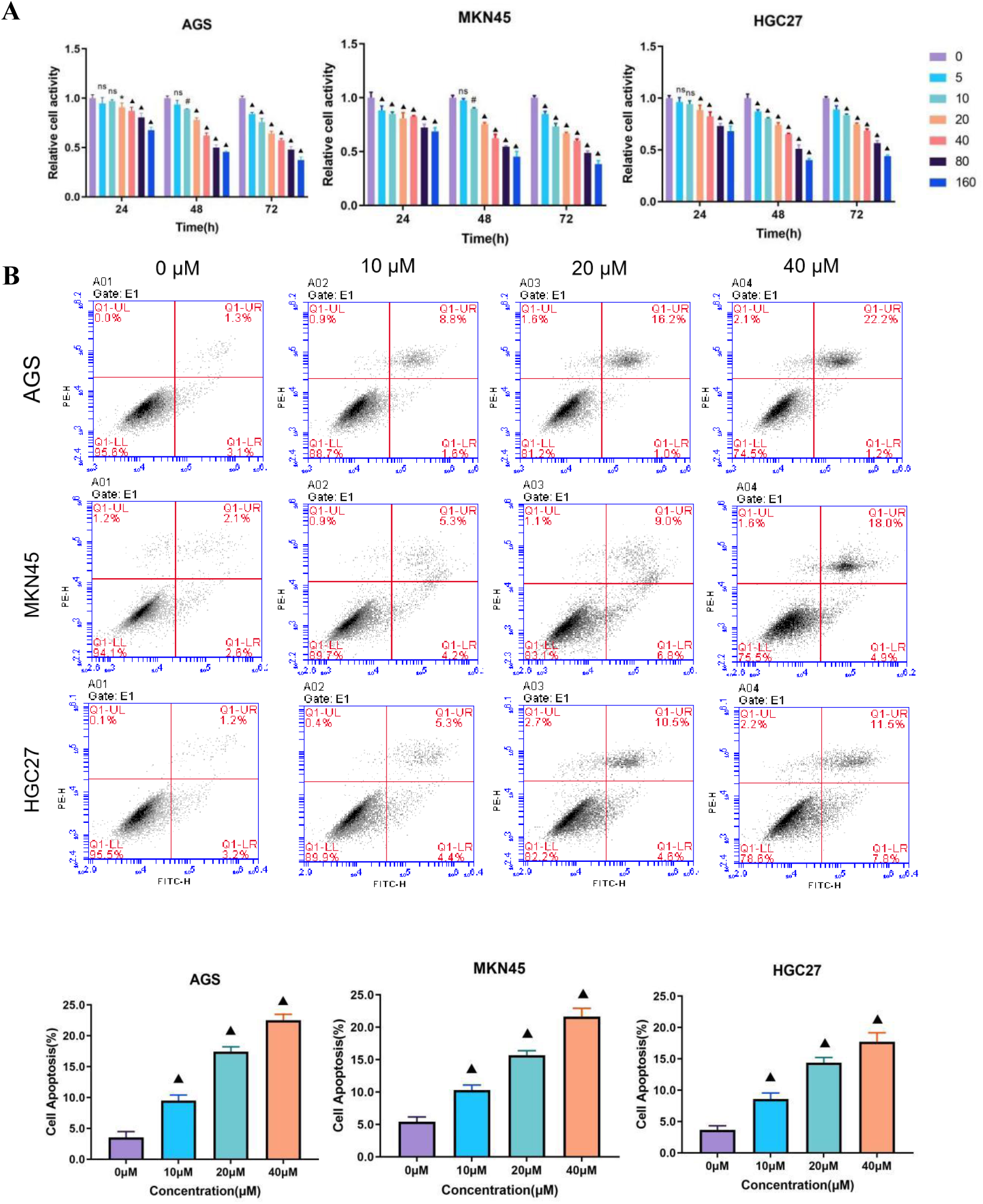
Effect of OXA on the proliferation and apoptosis of GC cells. (A) AGS, MKN45, and HGC27 cells were treated with OXA at various concentrations (0, 5, 10, 20, 40, 80, and 160 μM) for 24, 48, and 72 hours. The proliferation of GC cells was assessed using the CCK-8 assay. (B) AGS, MKN45, and HGC 27 cells were treated with OXA at various concentrations (0, 10, 20, and 40 μM) for 48 hours. The apoptosis rates of GC cells were measured using flow cytometry with the Annexin-FITC/PI double staining method. Statistical significance is indicated as follows: “ns” indicates no statistically significant difference, “*” indicates *P* < 0.05, “#” indicates *P* < 0.01, and “▴” indicates *P* < 0.001.

#### 1.2 Oxaliplatin induces immunogenic cell death in gastric cancer cells

ICD is primarily mediated by DAMP, and involves surface-exposed CRT, ATP secretion, and HMGB1 release. First, we examined the effects of OXA on ICD in GC cells. We treated AGS, MKN45 and HGC27 cells at different concentrations (0,10,20 and 40 μM) of OXA for 48 h, and performing immunofluorescence experiments, the results showed that the obvious accumulation of CRT could be observed on the GC cell membrane, In the control group, CRT was mainly distributed in the cytoplasm (Figure 2A, Supplementary Figures S1 and S2). These results confirmed that OXA translocated CRT to the cell membrane in GC cells.

**Figure 2.**
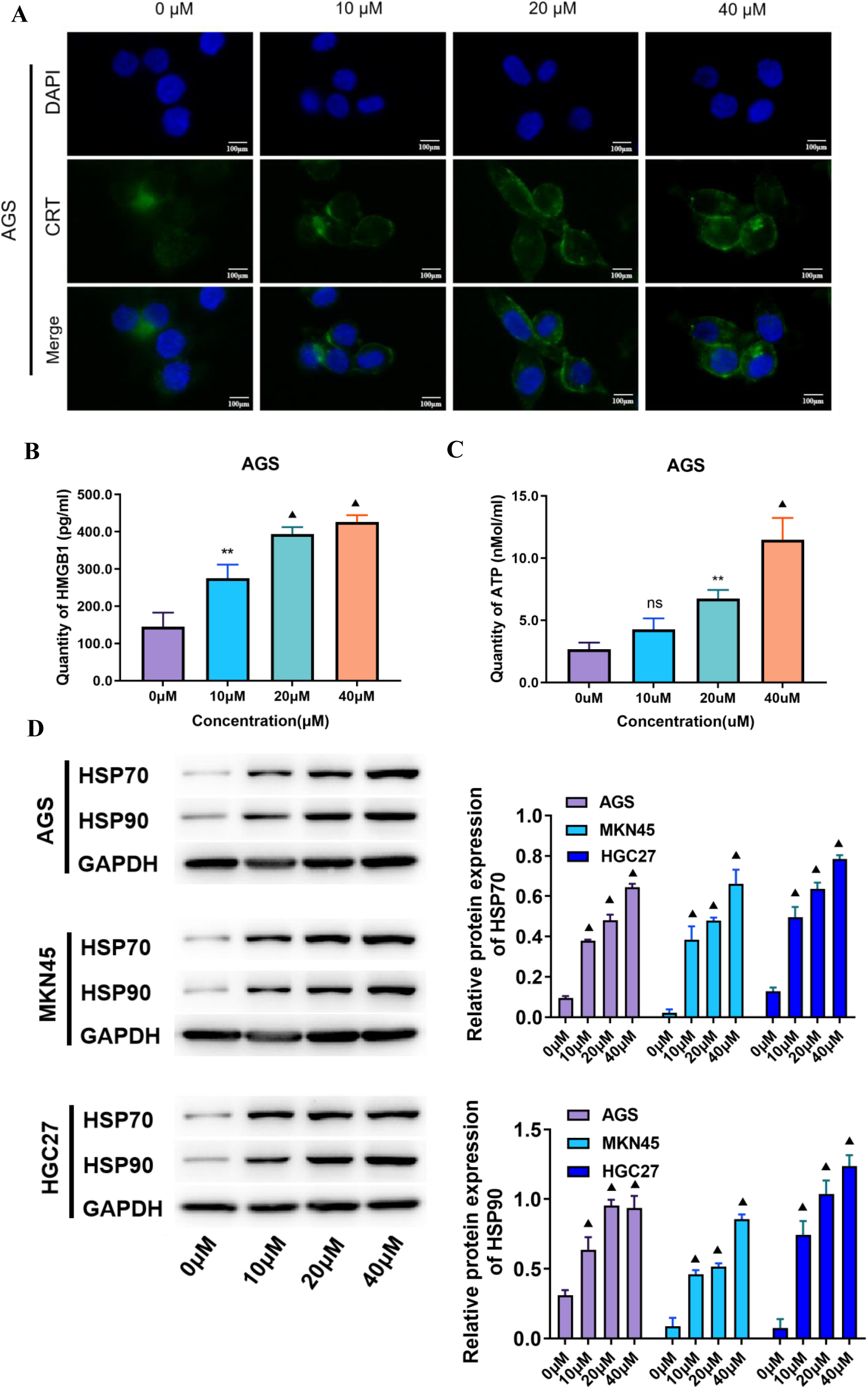
OXA induces Immunogenic Cell Death in GC cells. (A) AGS cells were treated with OXA at various concentrations (0, 10, 20, and 40 μM) for 48 hours. The changes in CRT were observed via immunofluorescence. (B) AGS cells were treated with OXA at various concentrations (0, 10, 20, and 40 μM) for 48 hours. The content of HMGB1 in the cell supernatant was measured using ELISA, and a statistical chart was generated.(C) AGS cells were treated with OXA at concentrations (0, 10, 20, and 40 μM) for 48 hours. The ATP content in the supernatant was quantified using ATP assay kit, and a statistical graph was created. (D) AGS, MKN45, and HGC27 cells were treated with OXA at various concentrations (0, 10, 20, and 40 μM) for 48 hours. Western blot analysis was performed to assess the expression levels of HSP70 and HSP90 proteins, followed by statistical analysis. Scale bar: 100 μm. Statistical significance is indicated as follows: “ns” indicates no statistically significant difference, “**” indicates *P* < 0.01, and “▴” indicates *P* < 0.001.

Next, to explore the effect of OXA on HMGB1 and ATP release in GC cells, we treated AGS, MKN45, and HGC27 cells at different concentrations (0,10,20 and 40 μM) for 48 h, and the cell supernatants were collected to detect the content of HMGB1 and ATP by ELISA. The results showed that the content of HMGB1 and APT was significantly higher than that of the control group, along with OXA concentration (*P* < 0.05, Figure 2 B, C and Supplementary Figure S1 and S2).

Furthermore, we also detected HSP70/90 protein expression in GC cells by western blotting. We treated AGS, MKN45, and HGC27 cells with different concentrations (0,10,20 and 40 μM) of OXA for 48 h, collected GC cells, and detected HSP70/90 protein expression by western blotting. The results showed that the 70/90 protein expression level in the other concentration groups was significantly higher than that in the control group (*P* <0.001, Figure 2 D).

### 2 OASL is able to inhibit oxaliplatin-induced immunogenic cell death in gastric cancer cells

#### 2.1 Effect of OASL on oxaliplatin-induced apoptosis in gastric cancer cells

To further clarify the relationship between OXA and OASL, we treated AGS, MKN 45, and HGC27 cells with different concentrations (0,10,20, and 40 μM) for 48 h, and OASL protein expression was measured by western blotting. The results showed that the protein expression level of OASL increased with an increase in OXA concentration compared to that in the control group (*P* <0.001, Figure 3A). Similarly, with the OASL mRNA expression level measured by RT-PCR, the increased OASL mRNA expression level and OXA concentration were statistically significant (*P* <0.05, Figure 3B). The results showed that the protein and mRNA expression levels of OASL increased with increasing OXA concentrations, suggesting that the expression level of OASL was closely related to OXA.

**Figure 3.**
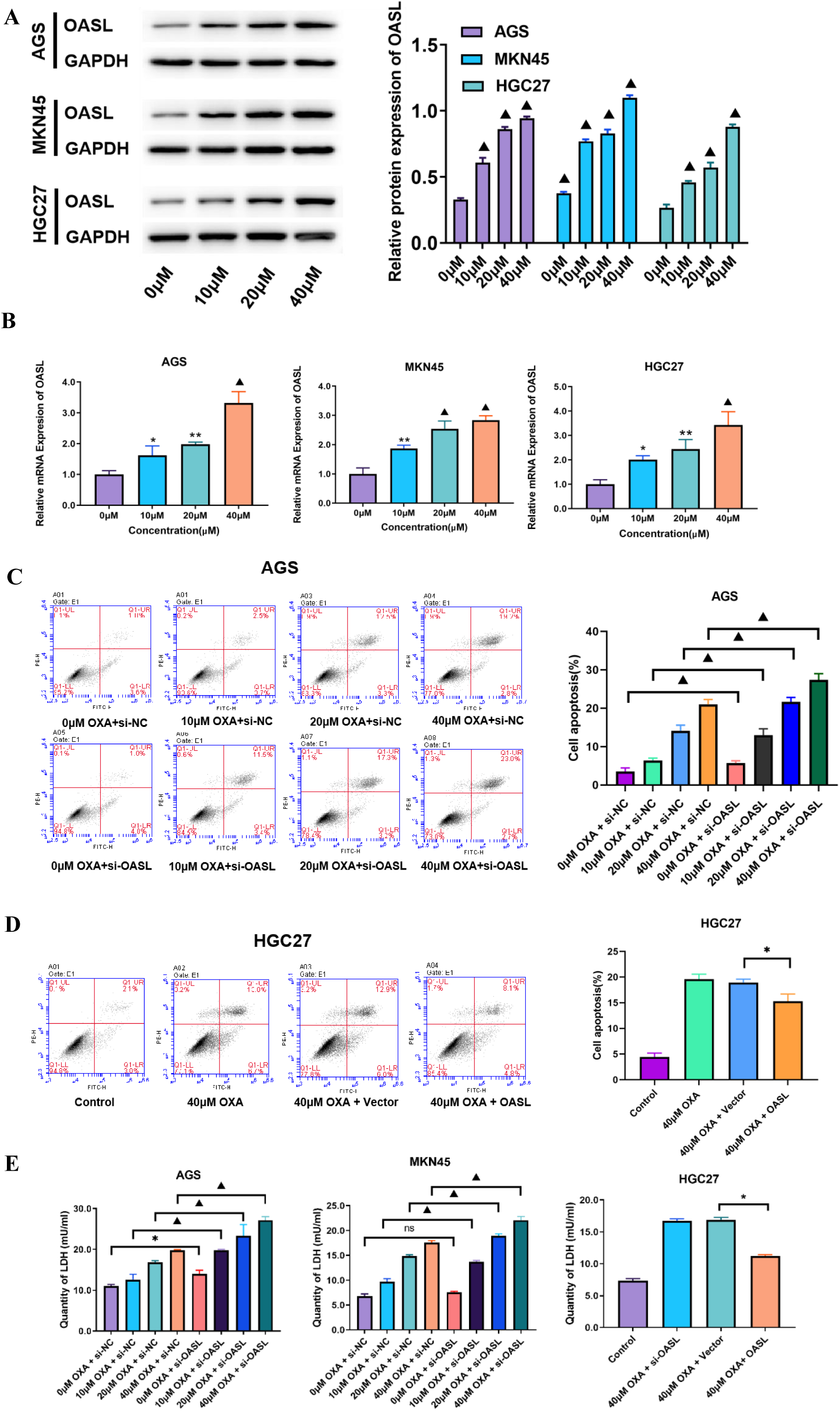
The Effect of OASL on Oxaliplatin-Induced apoptosis in GC cells. (A) AGS, MKN45, and HGC27 cells were treated with OXA at various concentrations (0, 10, 20, and 40 μM) for 48 hours. The Western blot analysis was performed to assess the protein expression levels of OASL in GC cells, followed by statistical analysis. (B) AGS, MKN45, and HGC27 cells were treated with OXA at various concentrations (0, 10, 20, and 40 μM) for 48 hours. RT-PCR was then utilized to measure OASL mRNA expression levels in the GC cells, followed by statistical analysis. (C) AGS cells underwent OASL knockdown and were subsequently treated with OXA at different concentrations (0, 10, 20, and 40 μM) for 48 hours. The apoptosis rate was measured by flow cytometry using the Annexin FITC/PI double staining method, and a statistical analysis of the apoptosis rate was conducted. (D) HGC27 cells were overexpressed with OASL and then treated with 40 μM OXA for 48 hours. Flow cytometry (using the Annexin FITC/PI double staining method) was employed to measure the apoptosis rate of AGS cells, along with the generation of a statistical chart reflecting the apoptosis rate. (E) Statistical graphs showed LDH content in AGS and MKN45 cells after OASL knockdown, followed by treatment with OXA at various concentrations (0, 10, 20, and 40 μM) for 48 hours, as well as in HGC27 cells after OASL overexpression followed by treatment with 40 μM OXA for 48 hours. The notation “ns” indicates no statistically significant difference, “*” indicates *P* < 0.05, “**” indicates *P* < 0.05, and “^▴^” indicates *P* < 0.001.

Next, we explored whether OASL affected the sensitivity of GC cells to OXA chemotherapy. AGS and MKN 45 cells were knocked down with OASL at different concentrations (0,10,20,and 40 μM) for 48 h, and the rate of apoptosis was determined using flow cytometry (Annexin-FITC/PI double staining). The results showed that in AGS and MKN45 cells, the apoptosis rate in the si-OASL group compared with that in the control group was statistically significant (*P* <0.05, Figure 3 C, Supplementary Figure S1). To further verify the effect on apoptosis, we tested the LDH content and showed that the si-OASL group had significantly more LDH content than the control group, with a statistically significant difference (*P* <0.05, Figure 3E). Similarly, we treated HGC27 cells with OASL and overexpressed it with 40 μM OXA for 48 h. Apoptosis and LDH levels in the 40 μM OXA + OASL group were significantly lower than those in the control group, showing a statistically significant difference (*P* <0.05, Figure 3D, E). Combined with the above results, it was concluded that OASL inhibited OXA-induced apoptosis in GC cells and reduced their chemosensitivity of GC cells to OXA.

#### 2.2 Effect of OASL on oxaliplatin-induced immunogenic cell death of gastric cancer cells

To further elucidate the effect of OASL on immunogenic death induced by oxaliplatin in gastric cancer cells, we examined the effect of OASL on ICD-associated DAMPs in OXA-induced GC cells in AGS and MKN45 cells. The experiments were divided into control, OXA, OXA+ si-NC, and OXA + si-OASL groups with an OXA concentration of 40 μM, and ICD-associated DAMPs were detected after 48h in AGS cells. The experimental results showed that the OXA + si-OASL group related DAMPs (CRT agglomeration on the membrane surface, the content of HMGB1 and ATP in the medium supernatant and HSP70/90 protein expression in cells) were significantly higher in the OXA + si-NC group (*P* <0.05, Figure 4 A-D). The same method was used for MKN45 cells (Supplementary Figure S4). Combined with the above results, we concluded that OASL knockdown enhances OXA induction of ICD in GC cells.

**Figure 4.**
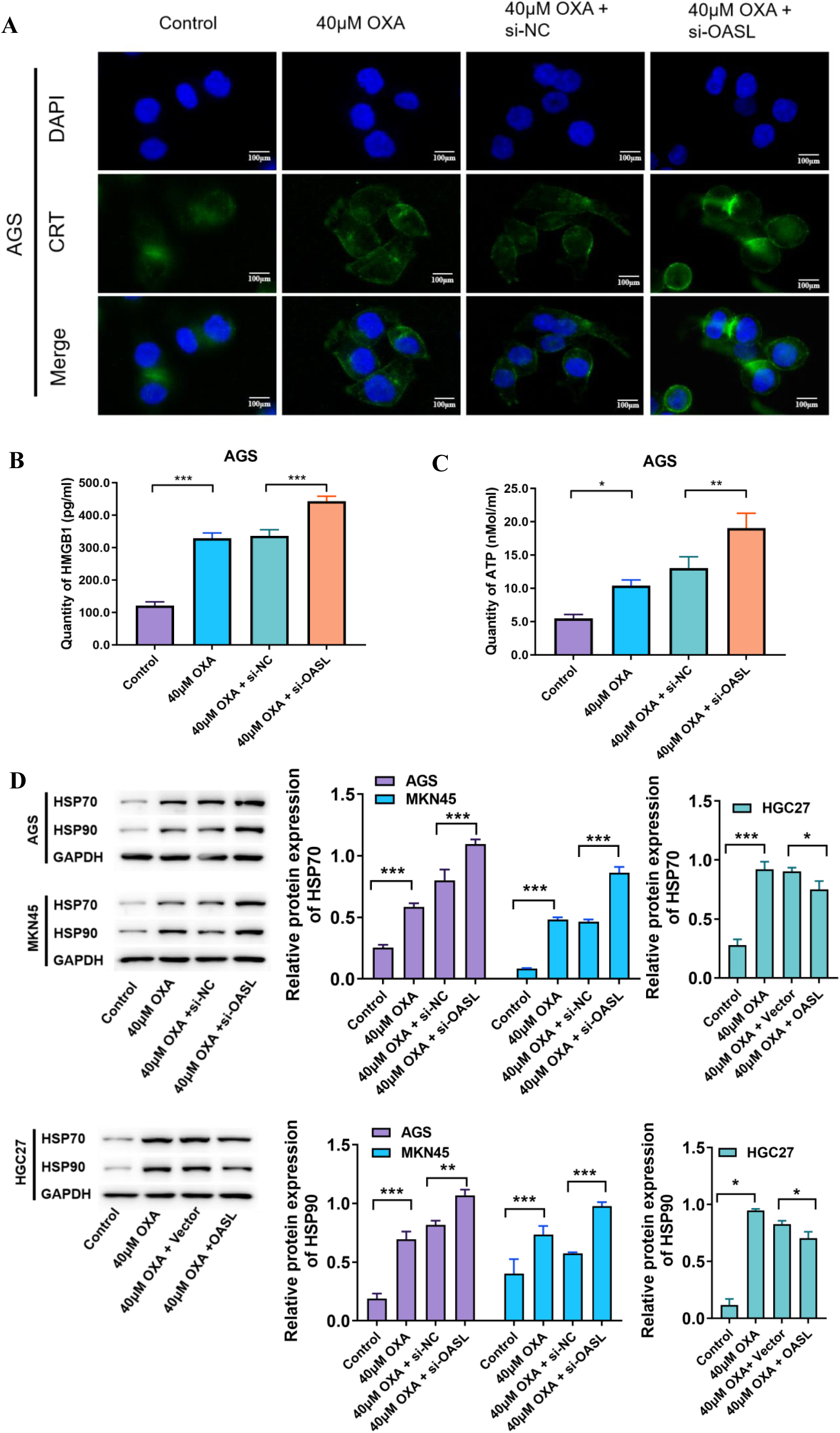
OASL can reduce immunogenic cell death induced by oxaliplatin in GC cells. (A) AGS cells were treated with 40 μM OXA combined with si-OASL for 48 hours, and changes in CRT were observed using immunofluorescence. (B) AGS cells were treated with 40 μM OXA combined with si-OASL for 48 hours, and the statistical plot of HMGB1 content in cell supernatants was detected using ELISA assay. (C) AGS cells were treated with 40 μM OXA combined with si-OASL for 48 hours, and statistical plot of ATP content in cell supernatant detected using ATP kit. (D) AGS and MKN45 cells were treated with 40 μM OXA in combination with si-OASL for 48 hours, and HGC27 cells were treated with 40 μM OXA in combination with overexpression of OASL for 48 hours. The expression levels of HSP70/90 protein in AGS, MKN45, and HGC27 cells were determined by Western blot, and statistical graphs were obtained. Scale bar: 100 μm. Statistical significance is indicated as follows: “ns” indicates no statistically significant difference, “*” indicates *P* < 0.05, “**” indicates *P* < 0.01, and “***” indicates *P* < 0.001.

To further validate the above results, we treated OASL overexpression combined with 40 μM OXA in HGC27 cells for 48 h, grouped them, and detected them as described above. The experimental results showed that the OXA + OASL group showed significantly less ICD related DAMPs (CRT on the surface of cell membrane, HMGB 1, and ATP content in the medium supernatant and HSP70/90 protein expression) (*P* <0.05, Supplementary Figure S5). Combined with the above results, it was concluded that the overexpression of OASL inhibited OXA to induce ICD in GC cells.

In conclusion, OASL reduced OXA levels to induce ICD in GC cells.

### 3 OASL regulation of the cGAS-STING pathway inhibits oxaliplatin-induced ICD

In previous experiments, we confirmed that OASL promoted the proliferation, invasion, and migration of GC cells and reduced the ICD of GC cells, thus reducing the chemotherapy sensitivity of OXA. However, the specific mechanism of action has not been elaborated on, and this section focuses on the following aspects.

#### 3.1 mRNA sequencing predicts the potential downstream signaling pathways of OASL

To explore the molecular mechanism of OASL in reducing ICD of GC cells, we first performed mRNA sequencing of MKN45 cells (OXA + si-NC group and OXA + si-OASL group) and visualized the mRNA expression levels of all detected (Figure 5 A); up-and down-regulated mRNA expression levels in each group using a volcano map (Figure 5B). To screen for differentially expressed mRNA, the screening criteria were set as differential multiples of 1. Reactome enrichment analysis is often used to detect the biological functions of differentially expressed genes. Reactome enrichment analysis of the differentially expressed genes showed significant enrichment in the second messenger signaling pathway (cGAMP) (Figure 5C). The cGAS recognizes and binds DNA in the cytoplasm, catalyzes the synthesis of ATP and GTP into cGAMP; cGAMP acts as a second messenger to bind and activate STING, promote the production of type I IFN and other inflammatory cytokines, and initiate the innate immune response. Based on the above findings, we speculated that the regulatory mechanism of ICD in GC cells induced by OASL is closely related to changes in the cGAS-STING signaling pathway.

**Figure 5.**
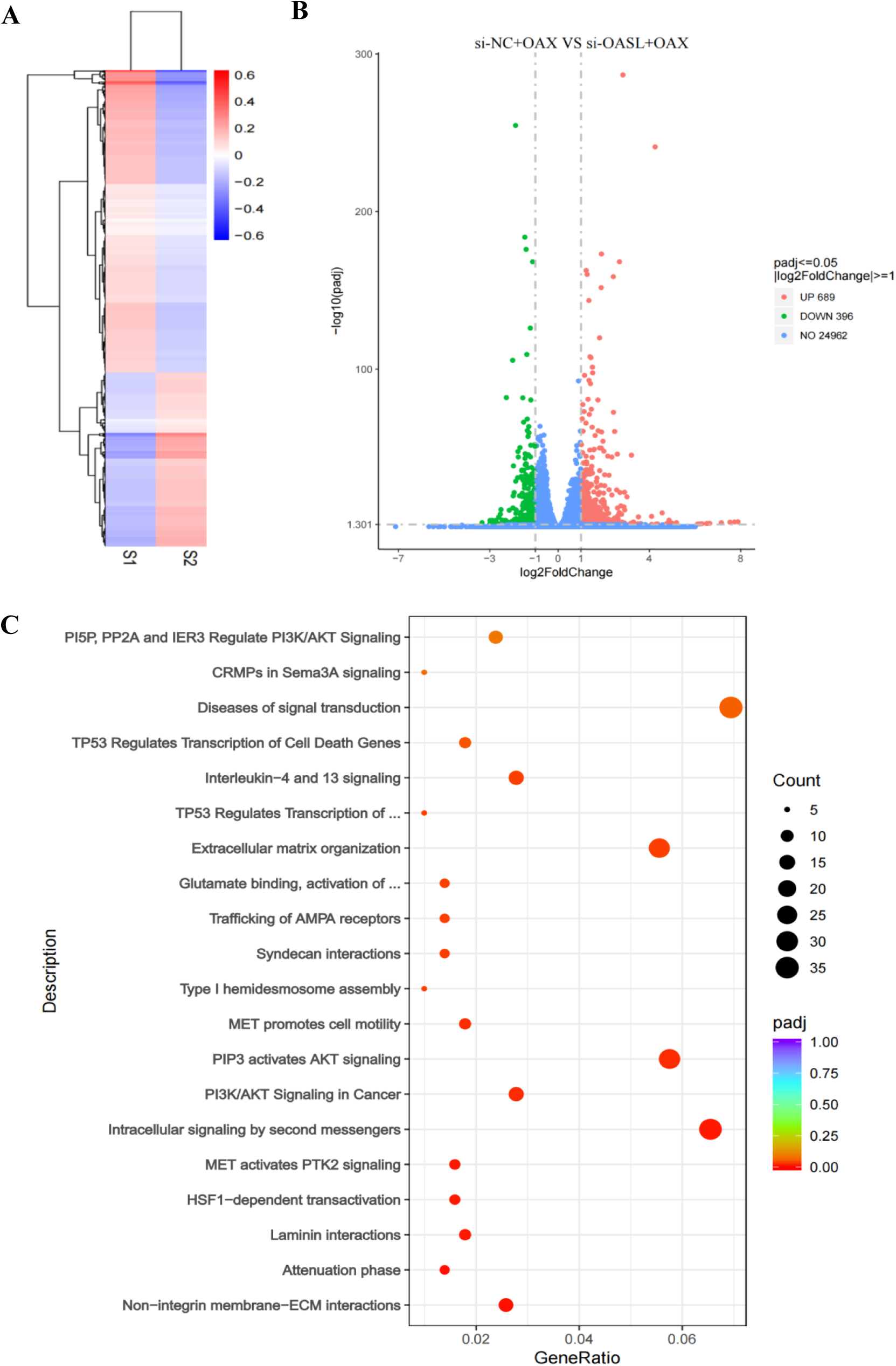
mRNA sequencing predicts potential downstream signaling pathways of OASL. (A) All the detected mRNA expression levels were visualized using the heatmap. (B) The Intervention up-regulated and down-regulated mRNA were displayed in groups using a volcano map. (C) Reactome enrichment analysis of the differential mRNA.

#### 3.2 OASL regulates the expression of key proteins of the cGAS-STING pathway

To further explore the relationship between OASL, cGAS, and STING, we first explored the correlation between OXA and the cGAS-STING signaling pathway. AGS, MKN45, and HGC27 cells were treated with OXA at various concentrations (0,10,20,and 40 μM) for 48 h, and western blotting was performed to detect the expression of key proteins of the cGAS-STING signaling pathway. The results showed that the protein expression of cGAS, STING and IRF3 increased at other concentrations compared with the control groups (0 μM) (*P* <0.05, Figure 6A). The results showed that OXA was able to enhance the activation of cGAS-STING signaling in GC cells, and the expression of related proteins, such as cGAS, STING, and RIF3, increased with an increase in OXA concentration in a dose-dependent manner.

**Figure 6.**
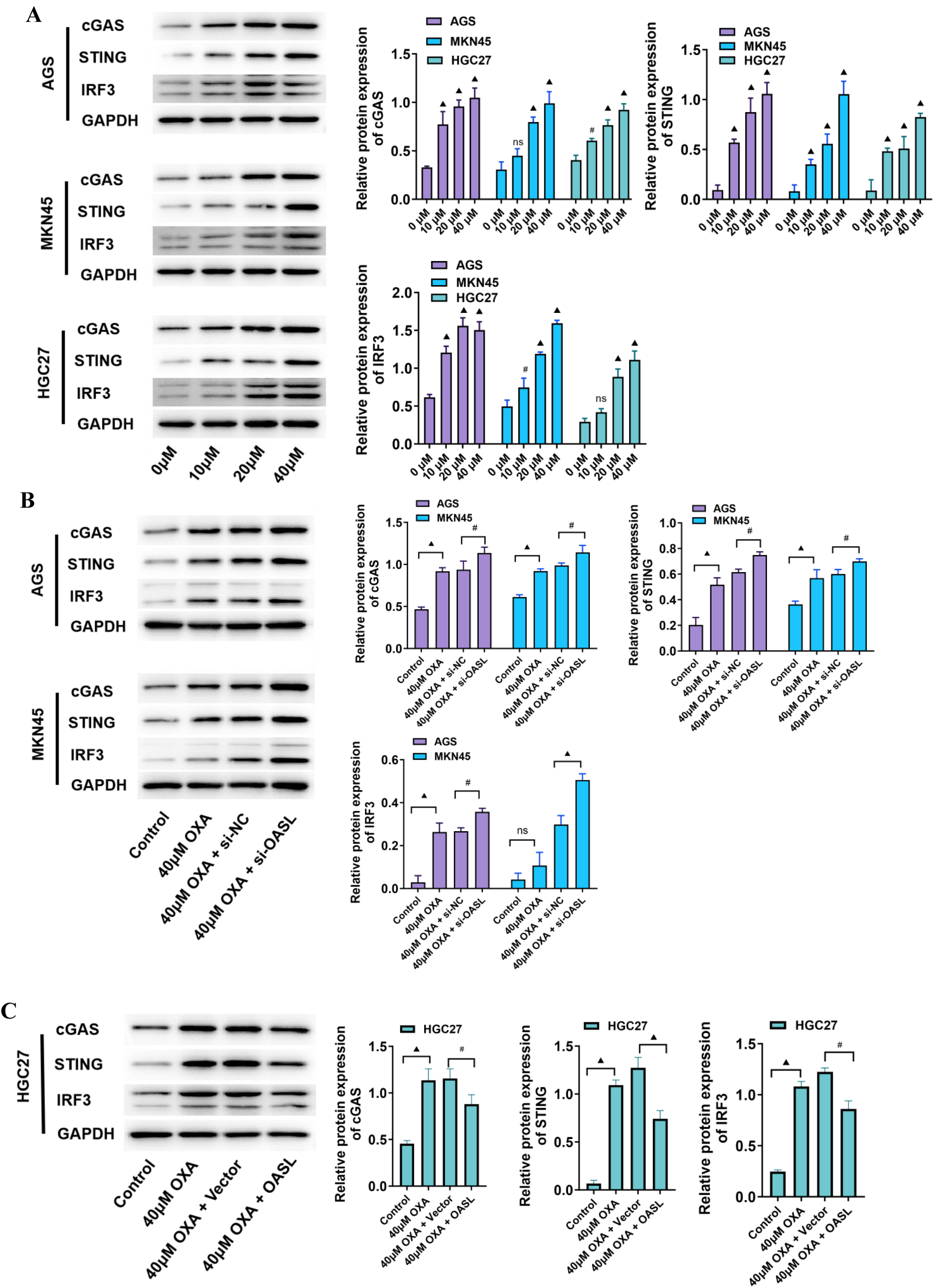
OASL regulates the expression of key proteins in the cGAS-STING pathway. (A) AGS, MKN 45, and HGC 27 cells were treated with OXA at various concentrations (0,10,20, and 40 μM) for 48 hours, and the expression of key proteins of the cGAS-STING signaling pathway was determined using Western blot.(B) AGS and MKN45 cells were treated with 40 μM OXA combined with si-OASL for 48 hours, and the expression of key proteins of the cGAS-STING signaling pathway was determined using Western blot. (C) HGC27 cells were treated with 40 μM OXA in combination with overexpression of OASL for 48 hours, and the expression of key proteins of the cGAS-STING signaling pathway was determined using Western blot. Statistical significance is indicated as follows: “ns” indicates no statistically significant difference, “#” indicates *P* < 0.05, and “▴” indicates *P* < 0.001.

Next, the relationship between OASL, cGAS, and STING was further analyzed to test the effect of OASL knockdown and overexpression on the protein expression levels of cGAS, SASNG, and RIF3 in GC cells. The experiments were divided into control, OXA, OXA + si-NC, and OXA + si-OASL groups, and 40 μM OXA was applied to AGS and MKN 45 cells for 48h, respectively, and the expression of key proteins of the cGAS-STING signaling pathway was determined by western blotting. The results showed that the protein expression of cGAS, STING and IRF3 in the OXA + si-OASL group increased (*P* <0.05, Figure 6B). HGC27 cells were treated with pcDNA3.1-OASL (OASL) combined with 40 μM OXA for 48 h and grouped and tested as described above. The results showed that the protein expression of cGAS, STING and IRF 3 was decreased in the OXA + OASL group than in the OXA + Vector group (*P* <0.05, Figure 6C). Based on these results, OASL can regulate the cGAS-STING pathway and exert biological effects.

#### 3.3 OASL relies on cGAS-STING signaling to regulate ICD due to oxaliplatin

To further verify that OASL can inhibit oxaliplatin-induced ICD through the regulation of the cGAS-STING signaling pathway, we designed a response experiment. The experiments were divided into control group, OXA group, OXA + si-NC group, OXA + si-OASL, OXA + OASL and OXA + si-OASL + inhibitor, and the detection of OXA concentration was 40 μ M, followed by AGS and MKN 45 cells (only key DAMPs such as HMGB 1 and HSP70/90 were detected in this section). In the AGS and MKN 45 cells, result display, Compared with the OXA + si-OASL group, After addition of cGAS-STING signaling pathway, Found that the reduction of HMGB1 in the supernatant of cell culture decreased, Reducreduced intracellular HSP70/90 protein expression, The differences were all statistically significant (*P* <0.05, Figure 7A, C); In HGC27 cells, compared with the OXA + OASL group, plus with Compound 3, an activator of the cGAS-STING signaling pathway, found that elevated levels of HMGB1 in the supernatant of the cell culture medium, increased intracellular expression of HSP70/90 protein, the differences were all statistically significant (*P* <0.05, Figure 7B, D). Based on the above findings, OASL may rely on the cGAS-STING signaling pathway to regulate the ICD of GC cells caused by OXA.

**Figure 7.**
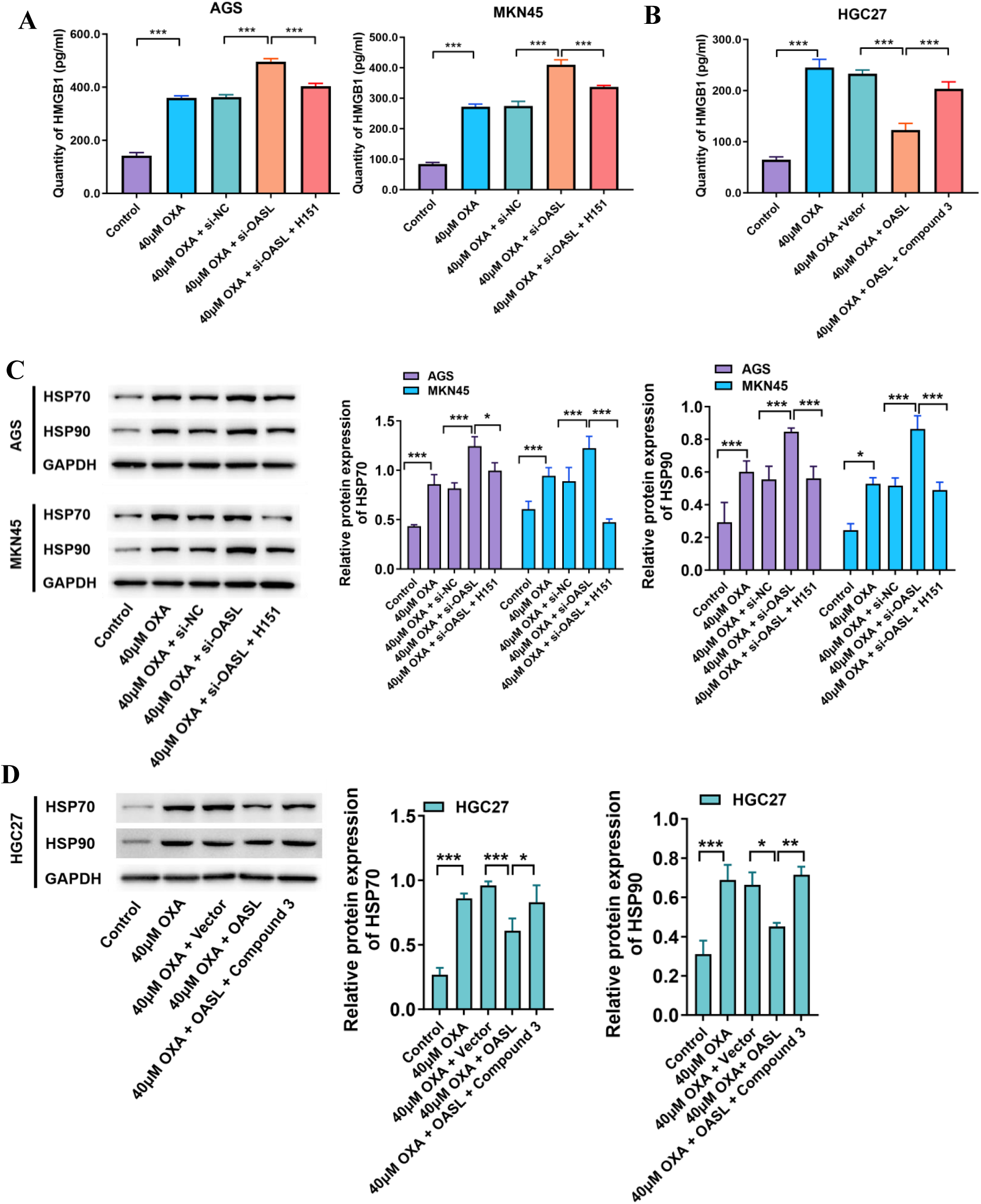
OASL-dependent cGAS-STING signaling to regulate oxaliplatin-induced ICD. (A)AGS and MKN 45 cells were treated with 40 μM OXA in combination with si-OASL and supplemented with H151(an inhibitor of the cGAS-STING signaling pathway) for 48 hours to determine the content of HMGB1 in the supernatant of cell culture medium.(B) HGC27 cells were treated with 40 μM OXA combined with overexpressed OASL and supplemented with Compound 3 (an activator of the cGAS-STING signaling pathway) for 48 hours to determine the content of HMGB1 in the supernatant of cell culture medium.(C) AGS and MKN 45 cells were treated with 40 μM OXA in combination with si-OASL and supplemented with H151(an inhibitor of the cGAS-STING signaling pathway) for 48 hours to determine HSP 70/90 protein expression using Western blot.(D) HGC27 cells were treated with 40 μM OXA combined with overexpressed OASL and supplemented with Compound 3 (an activator of the cGAS-STING signaling pathway) for 48 hours to determine HSP 70/90 protein expression using Western blot.Statistical significance is indicated as follows: “*” indicates P < 0.05, “**” indicates P < 0.01, and “***” indicates P < 0.001.

#### 3.4 OASL binds to cGAS proteins to inhibit the cGAS-STING signaling pathway

Based on the above results, we further explored the mechanism by which OASL regulates the cGAS-STING signaling pathway. The association between OASL and cGAS proteins was first demonstrated by the Co-IP results (Figure 8A). To further clarify the association between OASL and cGAS proteins, we administered OXA to the AGS and MKN cells. Immunofluorescence results showed an increase in OASL and cGAS expression in AGS and MKN cells in the OXA group compared with the control group, and the two could be directly combined (Figure 8B).

**Figure 8.**
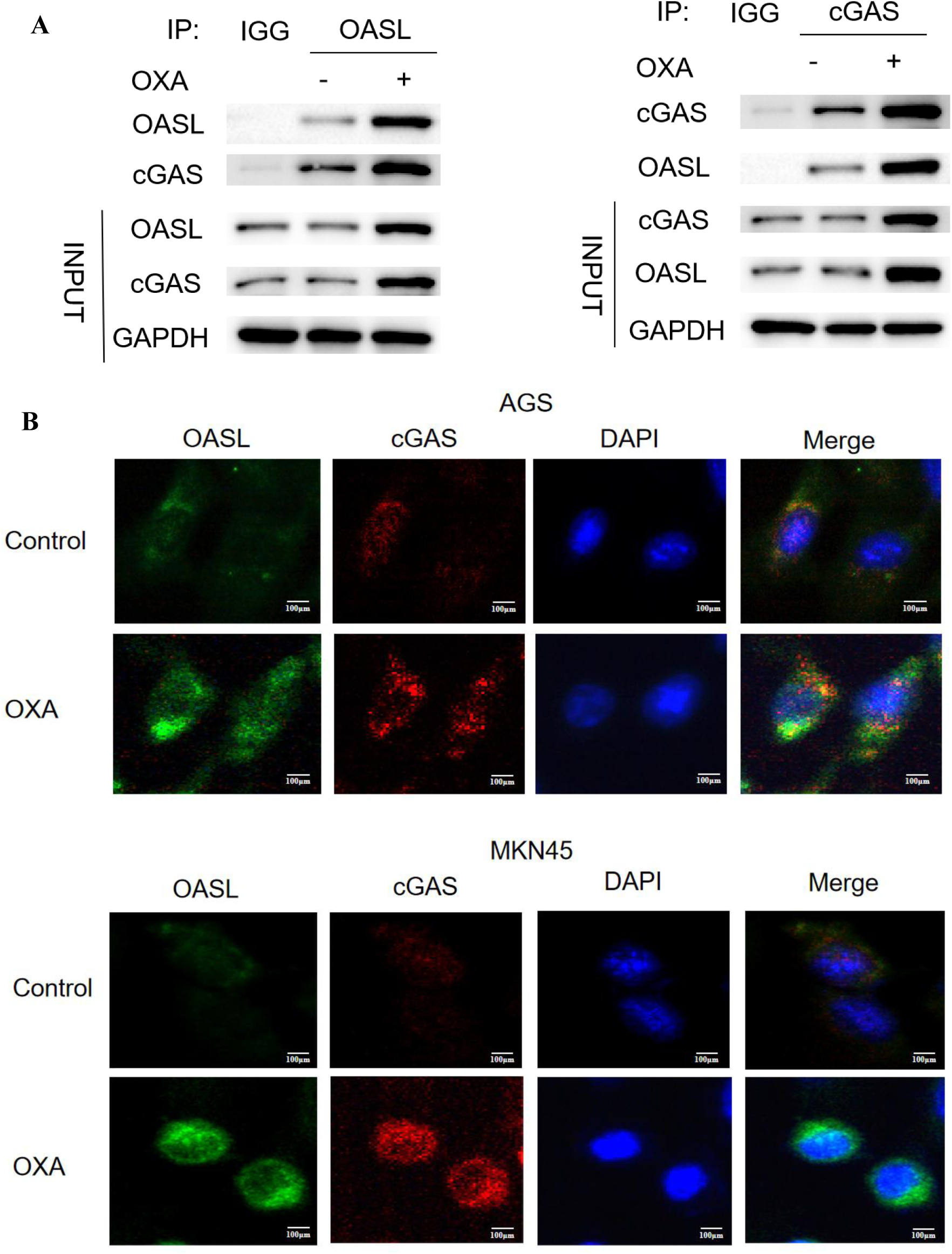
OASL binds to cGAS proteins to inhibit the cGAS-STING signaling pathway. (A) Co-IP results plots for OASL and cGAS.(B) Immunofluorescence plots verifying the binding of OASL and cGA by adding OXA treatment to AGS and MKN cells. Scale bar: 100 μm.

#### 3.5 Mouse tumor-bearing model verified that knockdown of OASL can inhibit ICD in gastric cancer cells caused by oxaliplatin

The above in vitro experiments demonstrated that**_10_**O**_0μm_**ASL can inhibit cGAS-STING signaling to reduce the ICD of GC cells. To further verify the mechanism of action of OASL in GC cells, we conducted in vivo experiments. C57 / BL6 mice were subcutaneously vaccinated with OASL shRNA knockdown MFC cells and NC-transfected MFC cells, and the experiments were divided into four groups: sh-NC, sh-OASL, OXA + sh-NC, and OXA + sh-OASL. The volume decreased in the sh-OASL group compared to the sh-NC group, and the OXA + Osh-OASL group was statistically significant (*P* <0.05, Figure 9A, B). The weight of the tumor-bearing tissue was significantly lower in the sh-OASL group than that in the sh-NC + OASL group (*P* <0.01, Figure 9C). The results showed that knockdown of OASL could inhibit the growth of MFC cells in mice and improve OXA chemotherapy sensitivity.

**Figure 9.**
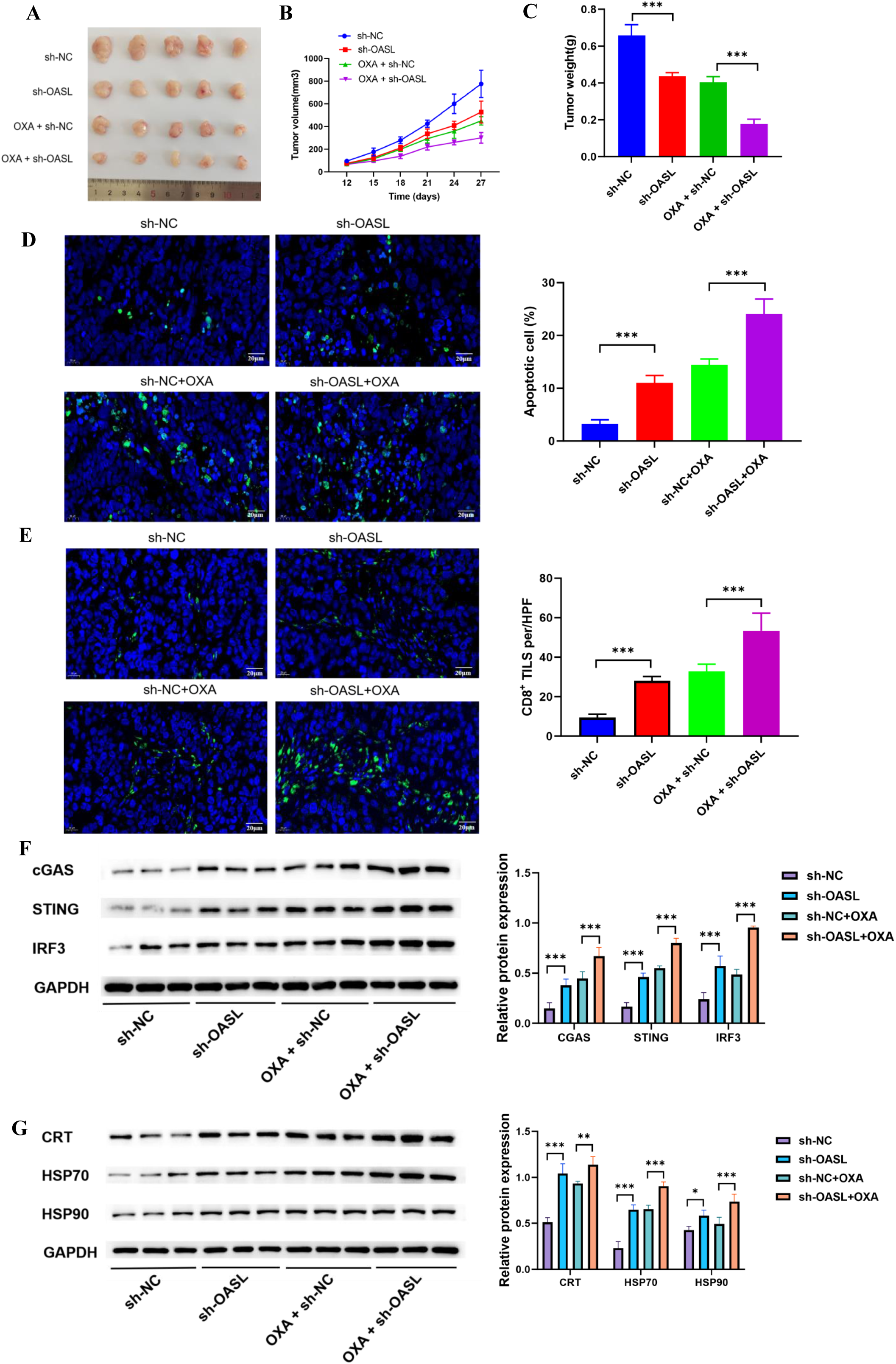
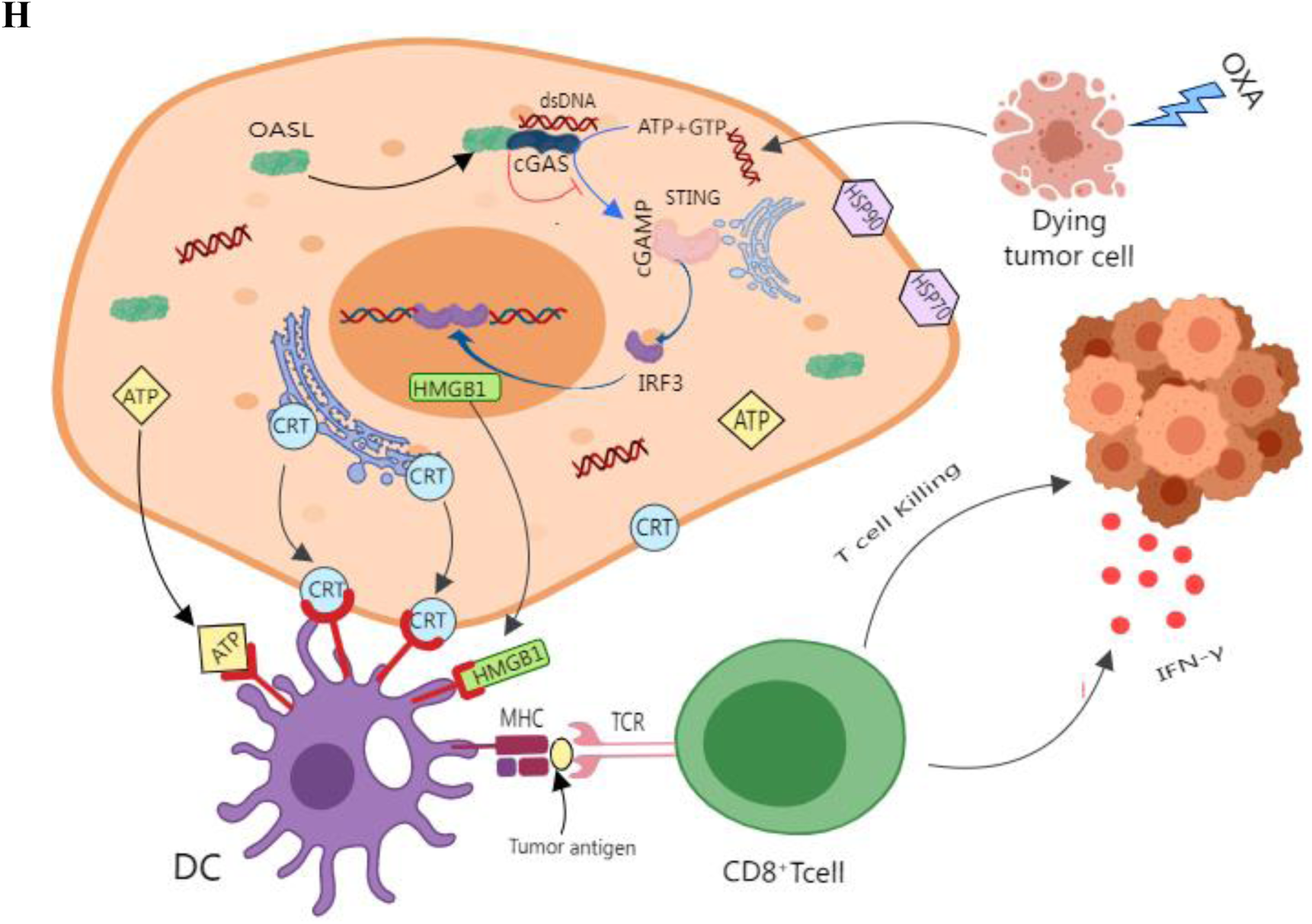
A mouse tumor-bearing model verified that knockdown of OASL improved OXA-induced ICD in GC cells. (A) In vivo tumorigenesis of MFC cells with sh-OASL in mice.(B) Vernier calipers measured the length and width of subcutaneous tumor-bearing tissue of inoculated mice from 12d after inoculation, and plot the growth curve of tumor-bearing tissue.(C) Weight statistical plot of the tumors.(D) Apoptosis and statistics of mouse tumor-bearing tumors detected by TUNEL staining.(E) CD8 expression and statistics of mouse tumor bearing by immunohistochemical fluorescence staining.(F) Expression and statistical plots of key proteins of cGAS-STING signaling pathway determined using Western blot.(G) Expression and statistical plots of the CRT and HSP 70/90 proteins associated with ICD were determined using Western blot. (H) The simulation diagram illustrating the mechanism how OASL exerts its biological effects. Scale bar: 20 μm. Statistical significance is indicated as follows: “*” indicates P < 0.05, “**” indicates P < 0.01, and “***” indicates P < 0.001.

TUNEL staining is a highly sensitive method for detecting apoptosis. The apoptosis rate was calculated by counting positive and negative cells, which showed significantly more positive cells and significantly higher apoptosis rates in the sh-OASL group than in the sh-NC group and in the OXA + sh-OXL group than in the OXA + sh-NC group (*P* <0.01, Figure 9D). To further define the CD8^+^ for the infiltration of TIL cells, we further performed immunofluorescence staining for CD8 and showed that CD8 in the sh-OASL group was compared with the sh-NC group^+^TIL cells; CD8 in OXA + sh-OASL group compared with OXA + sh-NC group^+^TIL cells was significant (*P* <0.01, Figure 9 E).

To further clarify the effect of OASL on the cGAS-STING signaling pathway, we examined the expression of key proteins of the cGAS-STING signaling pathway by western blotting, and the results showed that compared with the sh-NC group, the protein expression levels of cGAS, STING, and IRF 3 were significantly higher in the sh-OASL group than in the OXA + sh-NC group, and the protein expression levels of cGAS, STING, and IRF 3 were significantly higher in the OXA + sh-OASL group (*P* <0.001, Figure 9F).

To further clarify the effect of OASL on ICD, we detected the expression of ICD-related proteins by western blot, increasing the protein expression levels of CRT and HSP70/90 in the sh-OASL group (*P* <0.05) and CRT and HSP70/90 compared with the OXA + sh-NC group, respectively (*P* <0.001, Figure 9G).

In conclusion, the mouse tumor-bearing model verified that knockdown of OASL enhanced the activation of the cGAS-STING signaling pathway and enhanced the induction of ICD in GC cells by OXA, thus increasing the sensitivity of oxaliplatin chemotherapy.

## Discussion

GC is a common malignant tumor of the digestive tract that seriously affects human health. At present, GC treatment is mainly surgery, supplemented by radiotherapy and chemotherapy, in addition to targeted and immune comprehensive therapy. At present, the CSCO recommends first-line therapy for advanced GC, with chemotherapy as the cornerstone, which can be combined with immunotherapy or targeted therapy according to different conditions. Therefore, chemotherapy is indispensable for treating advanced GC^[7]^. OXA is one of the most commonly used platinum chemotherapy drugs. Its mechanism of action is the complexation of platinum atoms with DNA, which inhibits the replication and transcription of tumor cells. However, in recent years, OXA has been found in clinical medications, and this process is often easy, leading to serious drug resistance^[31]^. Although chemotherapy can effectively retreat or temporarily eliminate tumors in the short term, long-term use of chemotherapeutic drugs in patients often leads to MDR, which is one of the main causes of treatment failure in GC patients^[32,33]^. Therefore, reversing oxaliplatin resistance and improving chemotherapy sensitivity remain a challenge.

We innovatively started from the anti-tumor immunological mechanism and speculated that OASL may play an important role in regulating OXA chemotherapy sensitivity. To verify the specific regulatory mechanism, we first determined whether OXA could induce apoptosis in GC cells in vitro. Our results showed that OXA inhibited GC cell proliferation and promoted apoptosis. We further verified whether ICD is produced during GC cell apoptosis induced by OXA. The generation of ICD is highly dependent on a class of DAMPs, which involve cell surface-exposed CRT, release of HMGB, and secretion of ATP to activate and recruit antigen-presenting cells (such as DC cells), followed by activation of effector T cells to produce a specific anti-tumor immune response^[34]^. Our results confirmed that OXA translocated CRT to the membrane surface in GC cells and exposed it to HMGB1 and ATP, with an increase in OXA concentration. The current study found that the immune response has a direct relationship with three DAMPs, including CRT, HMGB1, and ATP, and HSP70/90 acts as molecular chaperone^[35–37]^. HSP70/90, as a molecular chaperone, enables the presence of tumor-associated antigen peptides to MHC class I molecules on the surface of DC, which in turn can process DC and initiate a subsequent anti-tumor immune response. Bortezomib and anthracyclines can induce exposure of HSP70/90 to the cell surface and act as carriers for antigenic peptides^[38]^. Our results also confirmed that OXA significantly increased the level of HSP70/90 protein expression in GC cells, along with increased OXA concentration. In conclusion, OXA induces ICD in GC cells in a dose-dependent manner. Subsequently, we regulated the expression of OASL and treated cells with OXA to detect the expression or release of DAMPs. The results showed that knockdown of OASL enhanced the ICD of GC cells in OXA, whereas overexpression of OASL inhibited OXA-induced ICD in GC cells. In conclusion, OASL reduced the ICD of GC induced by OXA. This confirmed that OASL plays an important role in the maintenance of chemosensitivity in GC.

To explore the molecular mechanism by which OASL reduces OXA-induced ICD in GC cells, we first performed mRNA sequencing of MKN45 cells (si-NC + OXA and si-OASL + OXA groups). Through differential expression and reactome enrichment analysis, the results showed significant enrichment in the second messenger signaling pathway. In 2011, the Russell E. Vance et al found that STING itself directly senses c-di-GMP DNA sensor in animal cells^[39]^. Along with the research, it was not proposed until 2013 that cGAMP could directly activate STING as a second messenger, and cGAS was a key synthetic enzyme upstream of cGAMP^[40]^. At this point, the specific composition and activation process of the cGAS-STING signaling pathway are clearly shown; when the cytoplasm, free dsDNA binds with cGAS, promotes GTP and ATP synthesis, cGAMP, and promotes STING activation, the latter is transported to perinuclear microbodies to form the TBK1-STING-IRF3 complex, and finally induces the production of type I IFN^[41,42]^. The cGAS-STING signaling pathway is the major pathway responsible for identifying the immune response to cytosolic DNA^[43]^, With the current research, the cGAS-STING signaling pathway is also playing an increasingly important role in anti-tumor immunity. Based on the above, we hypothesized that OASL exerts its biological effects through cGAS-STING.

To further explore the relationship between OASL and cGAS-STING signaling, we first explored the correlation between OXA and cGAS-STING signaling, and the results showed that OXA could cause an increase in related proteins such as cGAS, STING, and RIF 3 in GC cells, with an increase in OXA concentration and dose dependence. Moreover, when OASL was knocked down, the protein expression of cGAS, STING and IRF3 increased in the OXA + si-OASL group compared with the OXA+ si-NC group; the opposite result was observed when OASL was overexpressed. Collectively, these results suggest that OASL regulates the cGAS-STING pathway to exert its biological effects. To further verify that OASL can inhibit ICD induced by OXA, which is achieved by regulating the cGAS-STING signaling pathway, we designed a response experiment. In AGS and MKN45 cells, the addition of the cGAS-STING signaling H151 inhibitor decreased the amount of HMGB1 and intracellular HSP70/90 protein compared to the OXA + si-OASL group. Similarly, the addition of cGAS-STING signaling pathway activator Compound 3 increased the amount of HMGB1 and intracellular HSP70/90 protein expression compared with the OXA+OASL group. Based on the above findings, OASL may rely on the cGAS-STING signaling pathway to regulate the ICD of GC cells caused by OXA. Based on the above results, we further explored the mechanism of OASL regulating cGAS, we confirmed the association between OASL and cGAS proteins by Co-IP and immunofluorescence, which is consistent with the report by Ghosh A et al., where OASL binds to the DNA sensor cGAS during DNA viral infection, thereby inhibiting IFN induction and enhancing DNA viral replication^[44]^。

A growing number of studies have confirmed that the infiltration of tumor lymphocytes (tumor-invasive lymphocytes, TILs) is closely related to the overall survival of GC patients, and that high levels of tumor lymphocytes can significantly reduce the risk of recurrence and mortality in patients^[45–47]^. As the major component of TILs, CD8^+^TIL are considered to be one of the main drivers of anti-tumor immunity^[48]^. In the immune checkpoint treatment of GC, CD8^+^ infiltration of TIL can be used as a potential biomarker for prognosis prediction. In this study, in vivo experiments showed that OASL knockdown and OXA treatment significantly activated the cGAS-STING signaling pathway and enhanced the ICD effect, whereas knockdown of OASL promoted CD8^+^ infiltration of TIL cells. However, after the addition of OAX and CD8^+^ cells, the significantly increased infiltration of TIL cells was possibly associated with the enhanced ICD effect after the addition of OXA. Based on the above results, OXA effectively induced ICD in a mouse tumor-bearing model, and the combined knockdown enhanced OASL.

In conclusion, OASL was able to inhibit cGAS-STING signaling to reduce OXA-induced ICD in GC cells, leading to reduced sensitivity of GC cells to OXA chemotherapy.

## Data Availability Statement

The data that support the findings of this study are available on request from the corresponding author. The data are not publicly available due to privacy or ethical restrictions.

## Acknowledgement

All authors have agreed to the publication of this manuscript.

## Ethics Statement

The study protocol was approved by the ethics committee of Shandong Cancer Institute for Human Study and was conducted according to the principles of the Declaration of Helsinki.

## Patient consent for publication

Not applicable

## Author Contributions

All authors made a significant contributions to the reported work, including in the conception, study design, experimental implementation, execution and/or interpretation; participated in drafting, revising or critically reviewing the article; gave final approval of the version to be published; agreed on the journal to which the article has been submitted; and agreed to be accountable for all aspects of the work.

Weizhu Zhao and Lugang Liu is responsible for experimental implementation;Yi Liu is responsible for data compilation and statistics; Haiying Yang is responsible for collating data; Longgang Wang and Jie Chai are responsible for IHC; Lingling Zhang and Dong Sun are responsible for experimental design and technical guidance.

---

**Supplementary Figure S1.**
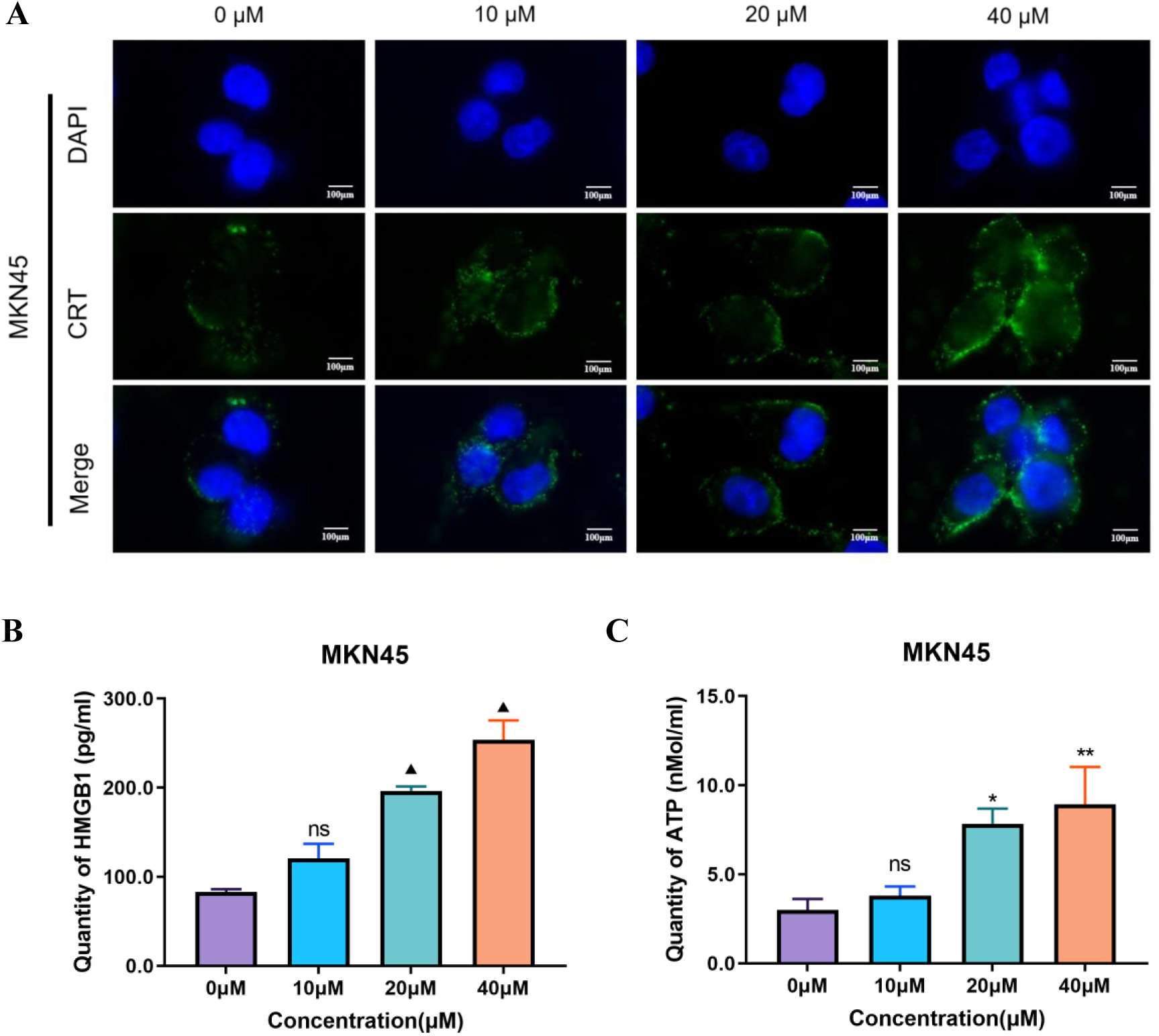
Oxaliplatin was able to induce immunogenic cell death in MKN45 cells. (A) MKN45 cells were treated with OXA at various concentrations (0,10,20 and 40 μM) for 48 hours, the changes in CRT were observed via immunofluorescence. (B) MKN45 cells were treated with OXA at various concentrations (0,10,20 and 40 μM) for 48 hours, the content of HMGB1 in the cell supernatant was measured using ELISA, and a statistical chart was generated.(C) AGS cells were treated with OXA at concentrations (0, 10, 20, and 40 μM) for 48 hours, the ATP content in the supernatant was quantified using ATP assay kit, and a statistical graph was created. Scale bar: 100 μm. Statistical significance is indicated as follows: “ns” indicates no statistically significant difference, “*” indicates *P* < 0.05, “**” indicates *P* < 0.01, and “▲” indicates *P* < 0.001.

**Supplementary Figure S2.**
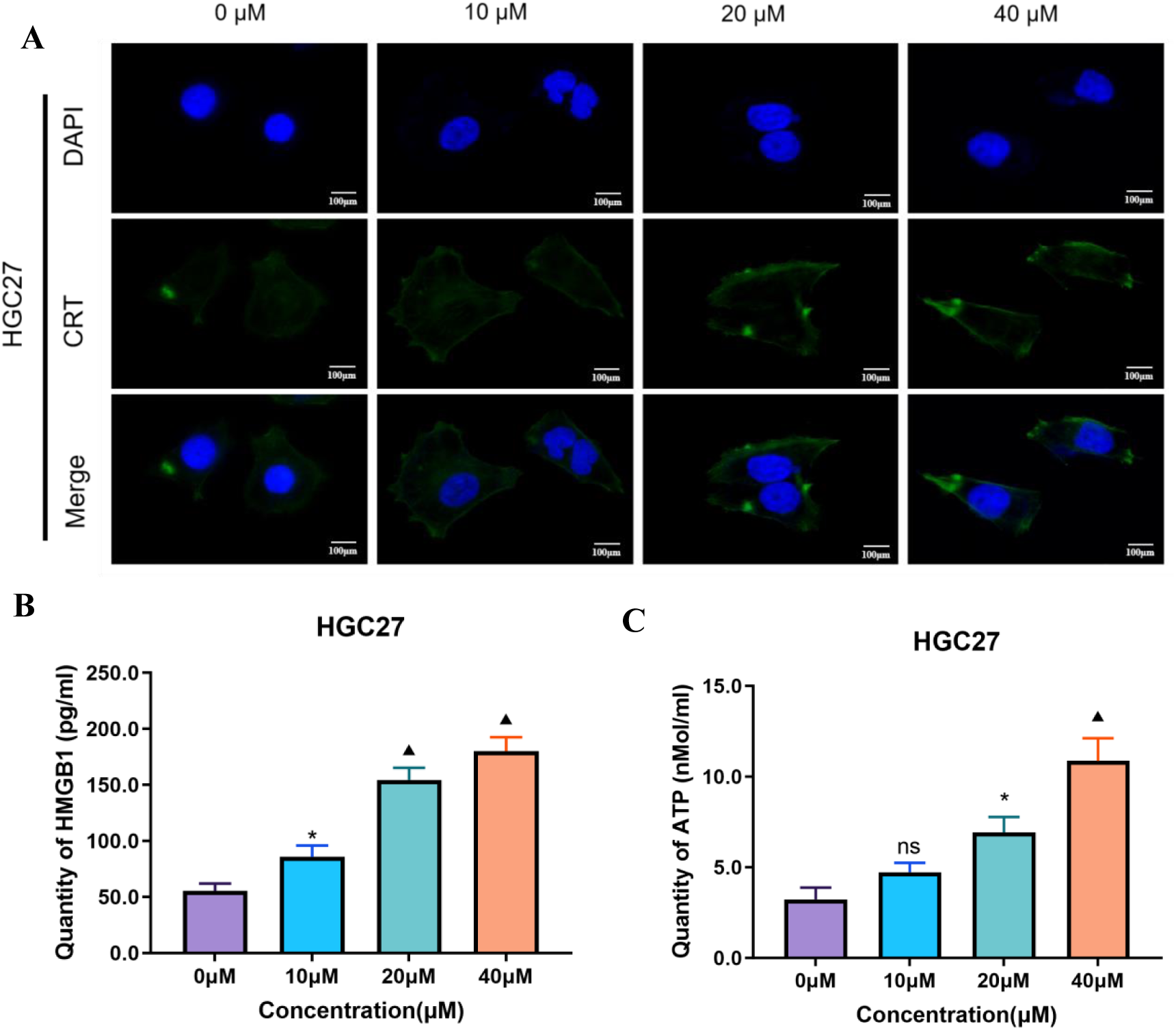
Oxaliplatin was able to induce immunogenic cell death in MKN45 cells. (A) HGC27 cells were treated with OXA at various concentrations (0,10,20 and 40 μM) for 48 hours, the changes in CRT were observed via immunofluorescence. (B) HGC27 cells were treated with OXA at various concentrations (0,10,20 and 40 μM) for 48 hours, the content of HMGB1 in the cell supernatant was measured using ELISA, and a statistical chart was generated.(C) AGS cells were treated with OXA at various concentrations (0, 10, 20, and 40 μM) for 48 hours, the ATP content in the supernatant was quantified using ATP assay kit, and a statistical graph was created. Scale bar: 100 μm. Statistical significance is indicated as follows: “ns” indicates no statistically significant difference, “*” indicates *P* < 0.05, and “▴” indicates *P* < 0.001.

**Supplementary Figure S3.**
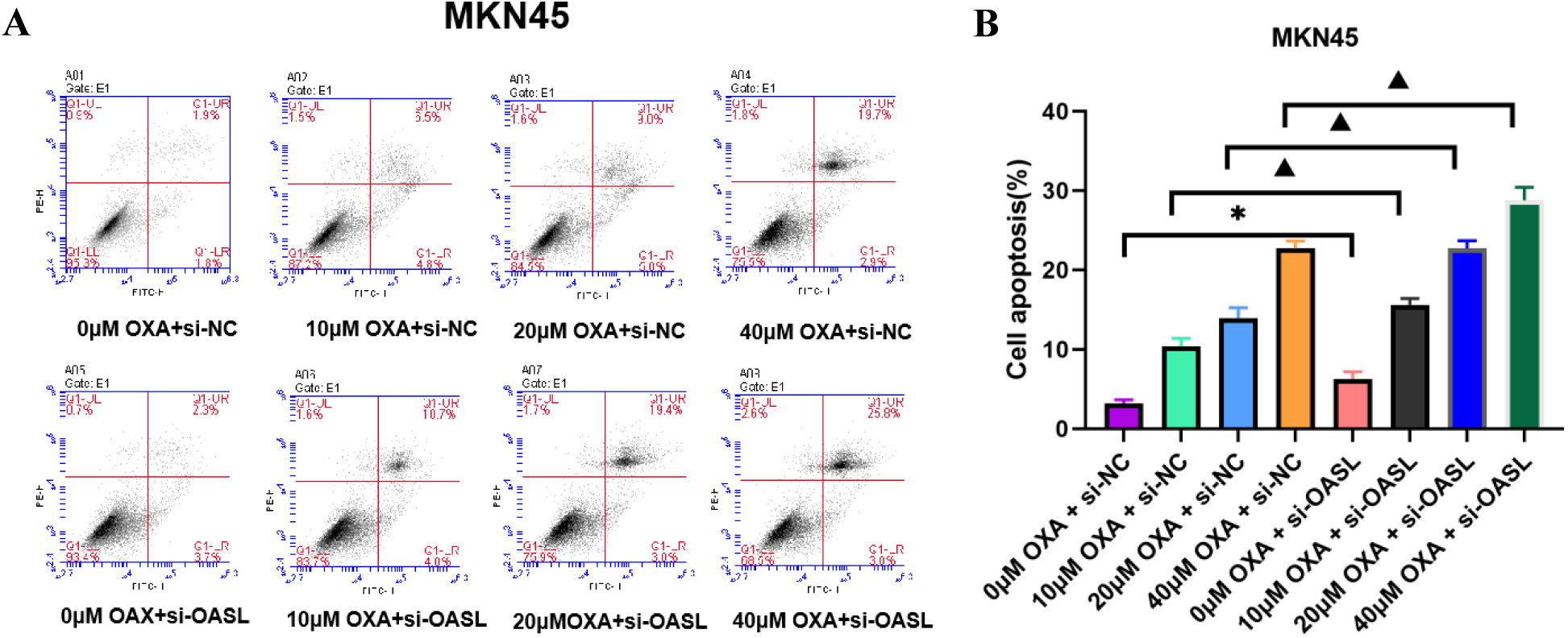
Effect of OXA combined with si-OASL on the apoptosis rate in MKN45 cells. MKN45 cells underwent OASL knockdown and were subsequently treated with OXA at various concentrations (0, 10, 20, and 40 μM) for 48 hours, the apoptosis rate was measured by flow cytometry using the Annexin FITC/PI double staining method, and a statistical analysis of the apoptosis rate was conducted. Statistical significance is indicated as follows: “*” indicates *P* < 0.05, and “▲” indicates *P* < 0.001.

**Supplementary Figure S4.**
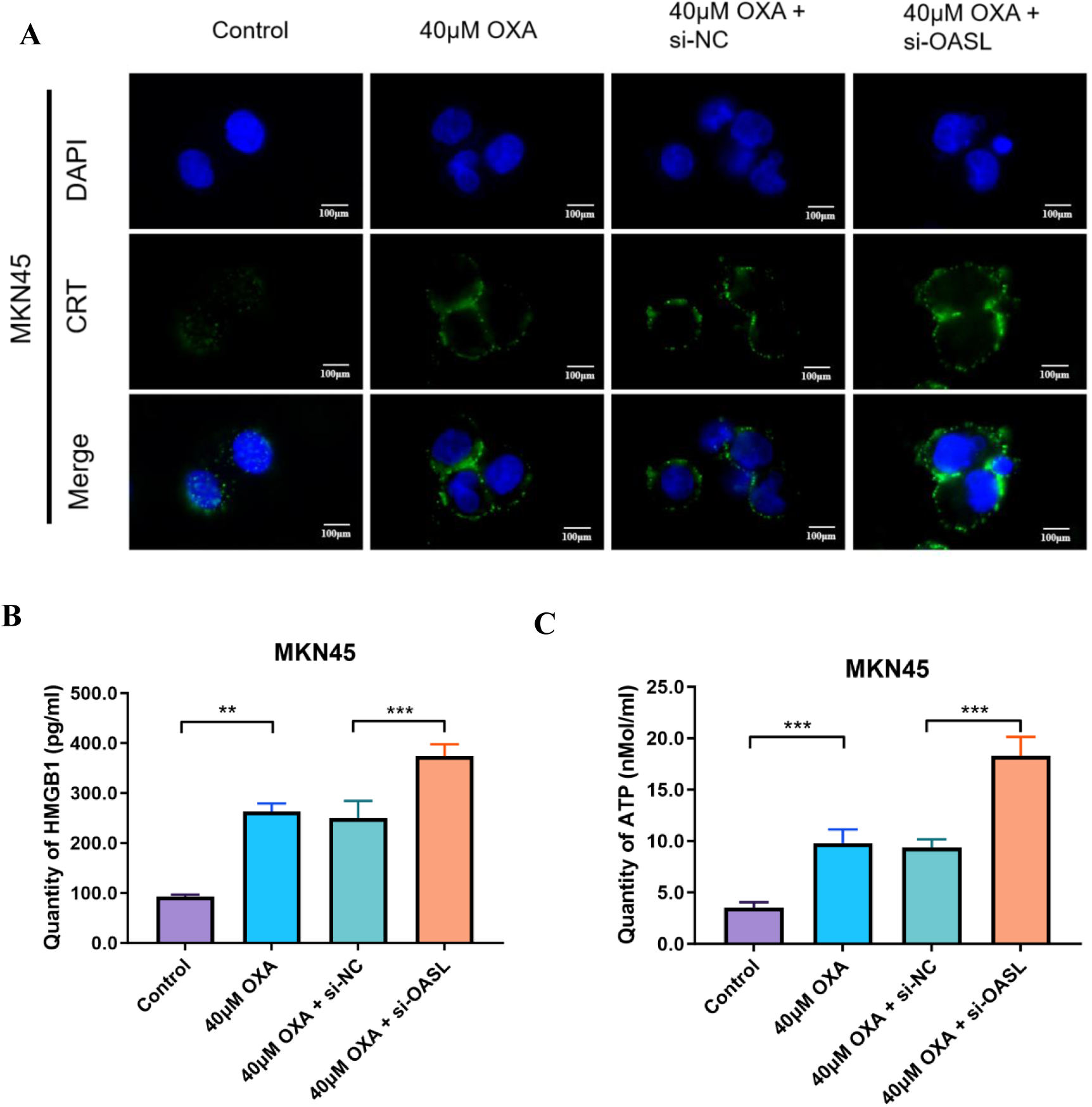
OASL was able to reduce oxaliplatin-induced immunogenic cell death in MKN45 cells. (A) MKN45 cells were treated with 40 μM OXA combined with si-OASL for 48 hours, and changes in CRT were observed using immunofluorescence. (B) MKN45 cells were treated with 40 μM OXA combined with si-OASL for 48 hours, and the statistical plot of HMGB1 content in cell supernatants was detected using ELISA assay. (C) MKN45 cells were treated with 40 μM OXA combined with si-OASL for 48 hours, and statistical plot of ATP content in cell supernatant detected using ATP kit. Scale bar: 100 μm. Statistical significance is indicated as follows: “**” indicates *P* < 0.01, and “***” indicates *P* < 0.001.

**Supplementary Figure S5.**
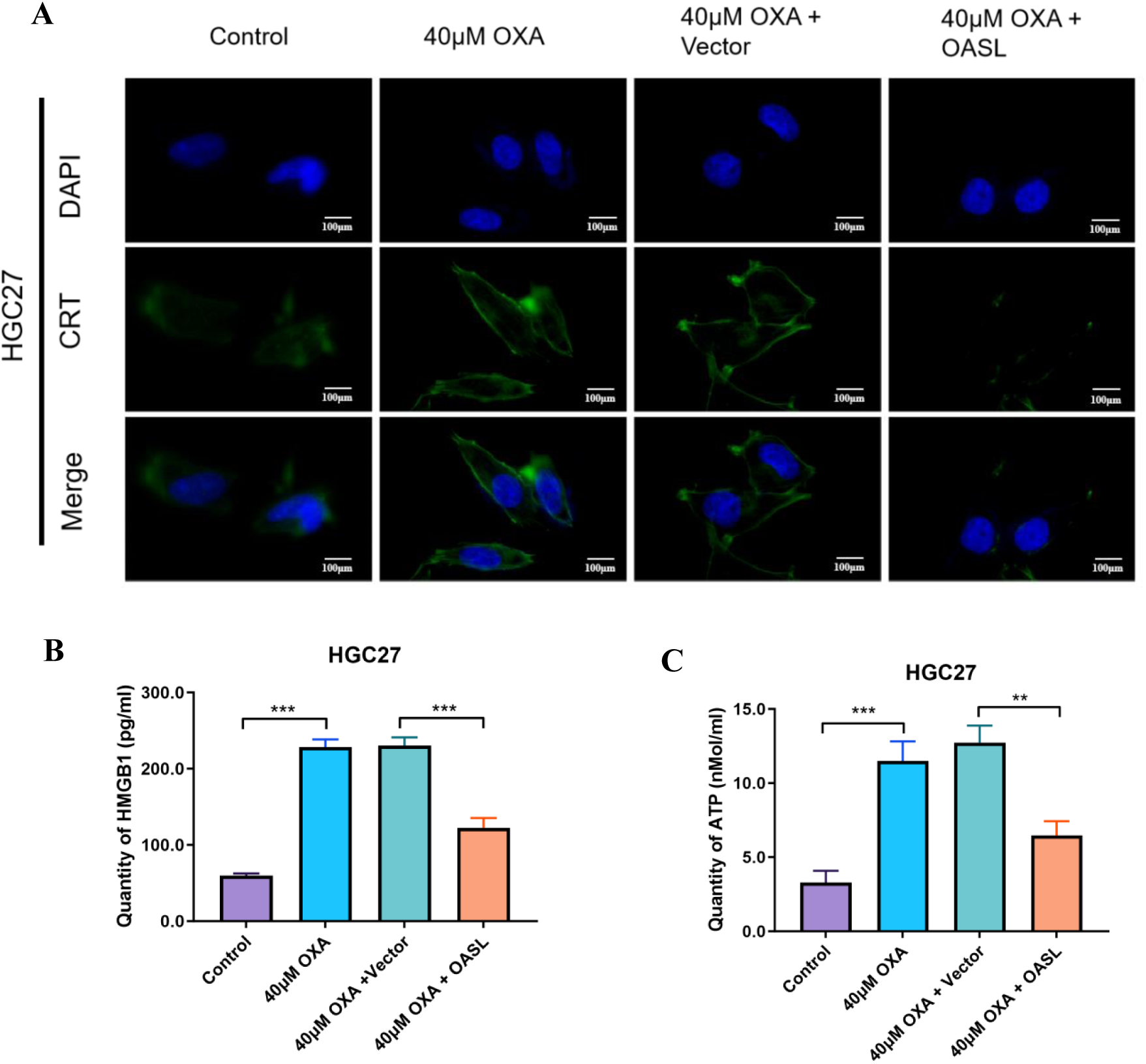
OASL was able to reduce oxaliplatin-induced immunogenic cell death in HGC27 cells. (A) HGC27 cells were treated with 40 μM OXA combined with overexpressed OASL for 48 h, and the changes in CRT were observed using immunofluorescence. (B) HGC27 cells were treated with 40 μM OXA combined with overexpressed OASL for 48 hours, and the statistical plot of HMGB1 content in cell supernatants was detected using ELISA assay. (C) HGC27 cells were treated with 40 μM OXA combined with overexpressed OASL for 48 hours, and statistical plot of ATP content in cell supernatant detected using ATP kit.

